# Resurrection of a Global, Metagenomically Defined Microvirus

**DOI:** 10.1101/743518

**Authors:** Paul C. Kirchberger, Howard Ochman

## Abstract

The *Gokushovirinae* (family *Microviridae*) are a group of single-stranded, circular DNA bacteriophages that have been detected in metagenomic datasets from every ecosystem on the planet. Despite their abundance, little is known about their biology or their bacterial hosts: isolates are exceedingly rare, known only from a very small number of obligate intracellular bacteria. By synthesizing circularized phage genomes from prophages embedded in diverse enteric bacteria, we produced viable gokushovirus phage particles that could reliably infect *E. coli*, thereby allowing experimental analysis of its life cycle and growth characteristics. Revived phages integrate into host genomes by hijacking a phylogenetically conserved chromosome-dimer resolution system, in a manner reminiscent of cholera phage CTX. Sequence motifs required for lysogeny are detectable in other metagenomically defined gokushoviruses, but we show that even partial motifs enable phages to persist in a state of pseudolysogeny by continuously producing viral progeny inside hosts without leading to collapse of their host culture. This ability to employ multiple, disparate survival strategies is likely key to the long-term persistence and global distribution of *Gokushovirinae*. The capacity to harness gokushoviruses as an experimentally tractable model system thus substantially changes our knowledge of the nature and biology of these ubiquitous phages.

## Introduction

Single-stranded circular DNA (ssDNA) phages of the family *Microviridae* are among the most common and rapidly evolving viruses present in the human gut (Lim *et al*., 2015; Minot *et al*., 2013). Within the *Microviridae*, members of the subfamily *Gokushovirinae* are regularly detected in metagenomic datasets from diverse environments, ranging from methane seeps to stromatolites, termite hindguts, freshwater bogs and the open ocean (Bryson *et al*., 2015; Desnues *et al*., 2008; Quaiser *et al*., 2015; Tikhe and Husseneder, 2018; Tucker *et al*., 2011).

Due to their small, circular genomes, full assembly of these phages from metagenomic data is easy (Creasy *et al*., 2018; Labonte and Suttle, 2013; Roux *et al*., 2012), and more than a thousand complete metagenome-assembled microvirus genomes have been deposited to public databases. In contrast, very few isolates of *Microviridae* exist, hardly representative of their diversity as a whole, and the only readily cultivable member of this family is *phiX*-174, which is classified to the subfamily *Bullavirinae* (Doore and Fane, 2016; Krupovic *et al*., 2016). While *phiX* and *phiX*-like phages are arguably the single most well-studied group of viruses, they are rare in nature and occupy a small specialist niche as a lytic predator of select strains of *Escherichia coli* (Michel *et al*., 2010). Conversely, the *Gokushovirinae* and several other (though not formally described) subfamilies of *Microviridae* are abundant in the environment but almost exclusively known from metagenomic datasets (Creasy *et al*., 2018; Szekely and Breitbart, 2016). Despite their prevalence in the environment, the only isolated gokushoviruses are lytic parasites recovered from the host-restricted intracellular bacteria, *Spiroplasma*, *Chlamydia* and *Bdellovibrio* (Brentlinger *et al*., 2002; Garner *et al*., 2004; Ricard *et al*., 1980). Given their regular occurrence in metagenomes from diverse habitats, it seems unlikely that *Gokushovirinae* only infect intracellular bacteria, and their lack of recovery from other hosts is puzzling.

Typical microviruses pack their 4–7 kilobase genomes, which encode 3–11 genes, into tailless icosahedral phage capsids (Roux *et al*., 2012). No microviruses encode an integrase, which has led to the assumption that they are lytic phages. However, the presence of prophages belonging to several undescribed groups of microviruses within the genomes of some *Bacteroidetes* and *Alphaproteobacteria* (Krupovic and Forterre, 2011; Zhan and Chen, 2018; Zheng *et al*., 2018) raises the possibility that some can be integrated through the use of host-proteins, in a similar manner to the XerC/XerD dependent integration of ssDNA *Inoviridae* (Krupovic and Forterre, 2015).

Most of the microvirus genomes assembled from metagenomic data lack multiple genes that are present in *phiX*-like phages (Roux *et al*., 2012). Markedly absent are: (*i*) a peptidoglycan synthesis inhibitor that leads to host cell lysis (although some phages appear to have horizontally acquired bacterial peptidases) and (*ii*) a major spike protein involved in host cell attachment (Doore and Fane, 2016; Roux *et al*., 2012). These proteins represent crucial elements in the infectious cycle of *phiX*, and their absence from other microviruses indicates that these phages might operate quite differently on a molecular level. By capturing a novel gokushovirus that is capable of lysogenizing enterobacteria, we characterized the strategies by which these viruses spread and persist in bacterial genomes, and demonstrate that they lead a decidedly different existence from that of previously characterized microviruses.

## Results

### A new, diverse group of gokushovirus prophages

Querying fully assembled bacterial genomes with the major capsid protein VP1 of gokushovirus *Chlamydia-*phage 4 returned 95 high-confidence hits (E-value < 0.0001) within the Enterobacteriaceae (91 from *Escherichia,* and one each from *Enterobacter*, *Salmonella*, *Citrobacter* and *Kosakonia*; **Table S1**). Although *Gokushovirinae* were previously known only as lytic phages, inspection of each of the associated genomic regions revealed the presence of integrated prophages 4300–4700 bp in length and having a conserved six-gene arrangement: VP4 (replication initiation protein), VP5 (switch from dsDNA to ssDNA replication protein), VP3 (scaffold protein), VP1 (major capsid protein), VP2 (minor capsid protein) and VP8 (putative DNA-binding protein. Most of the size and sequence variation in the prophages is confined to three regions: (*i*) near the C-terminus of VP2, (*ii*) in the non-coding region between VP8 and VP2, and (*iii*) within VP1, whose hypervariability is characteristic of the *Gokushovirinae* (Chipman *et al*., 1998; Diemer and Stedman, 2016) (**Figure 1A**).

**Figure 1.**
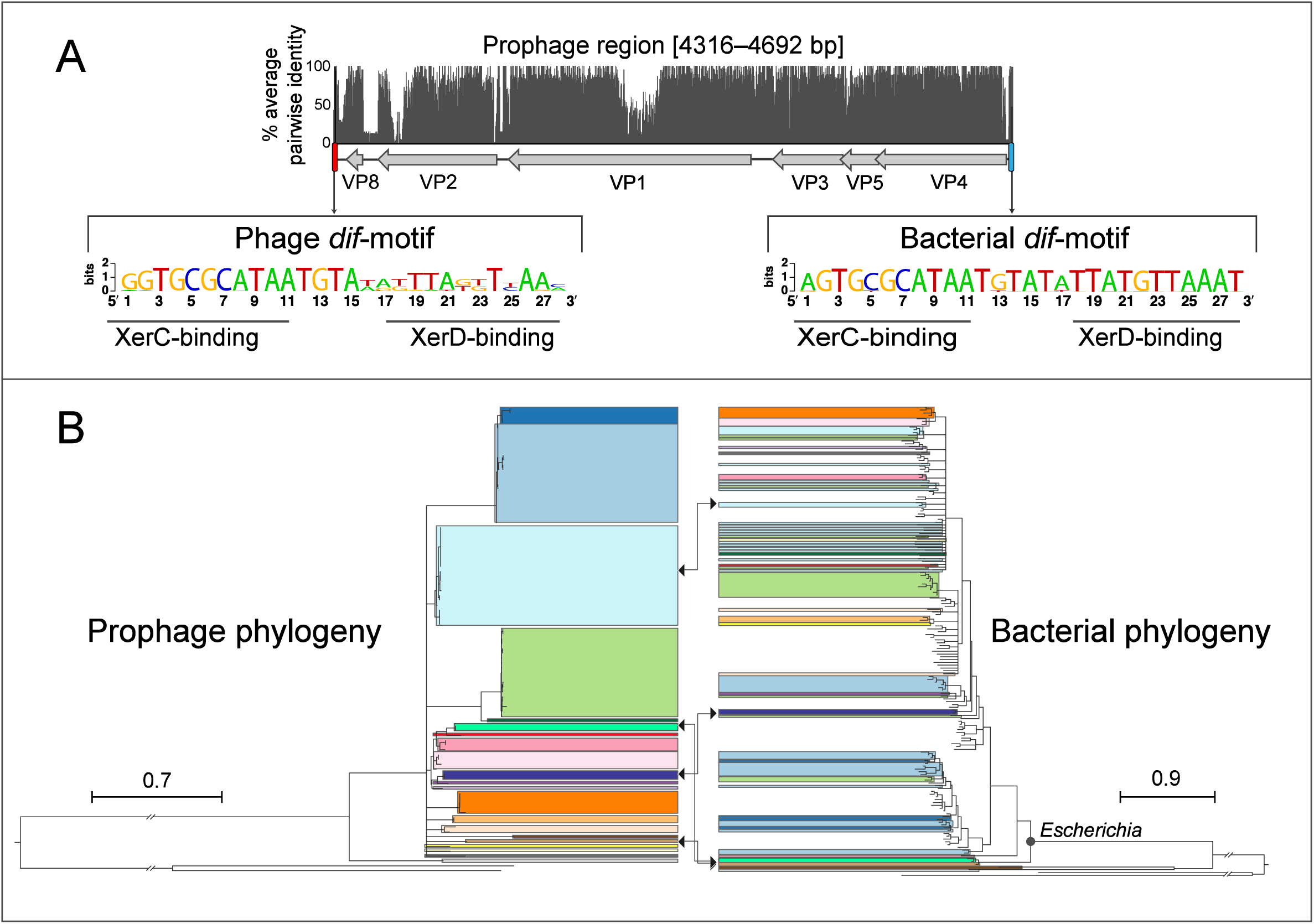
Gokushovirus prophages of Enterobacteria. (A) Genome organization and average pairwise nucleotide identities of gokushovirus prophages detected in *Escherichia*. Also shown are nucleotide sequence logos of the phage (downstream) and bacterial (upstream) *dif*-motifs with corresponding Xer-binding sites. (B) Phylogeny of gokushovirus prophages and their enterobacterial hosts. Phage clades with >95% average nucleotide identity are colored alike, and colors in the bacterial phylogeny correspond to the clade of their associated prophages found in each strain. Clades within the *Escherichia* phylogeny that are not colored represent ECOR collection strains, which are included to show the breadth of strain diversity. Clades with bootstrap support <70% are collapsed; arrows denote phages used for experimental analyses, as well as their corresponding hosts (see Methods). Ancestral branches were truncated by two legend lengths. Tree legend corresponds to substitutions/site. Accession numbers and details of prophages and corresponding hosts are listed in Tables S1 and S2.

All detected prophages are flanked by *dif*-motifs, which are 28-bp palindromic sites that are known to be the targets of passive integration by phages and other mobile elements (Blakely *et al*., 1993, Das *et al*. 2013). The *dif*-motifs upstream of the insertions are highly conserved and only differ by, at most, one nucleotide from the canonical *dif-motif* of *Escherichia coli.* These upstream *dif*-motifs consist of a central 6-bp spacer flanked by two 11-bp arms, which have previously been shown to bind tyrosine recombinases XerC/XerD during chromosome segregation and integration of mobile elements (Castillo *et al*. 2017). In contrast, the *dif*-motifs downstream of detected prophages are more variable, particularly in the spacer region and XerD-binding arms, representing the phage *dif*-motifs integrated along with the phage (**Figure 1A**).

A whole genome phylogeny of gokushovirus prophages shows a number of well-differentiated clades that each contains members with >95% average nucleotide identity (**Figure 1B**). Comparing the topology of the phage phylogeny with that of their *E. coli* host strains indicates that phage transfer and new insertions are common. Closely related bacteria can harbor dissimilar gokushovirus prophages, and, conversely, closely related prophages do not necessarily lysogenize only closely related *Escherichia* hosts. Despite the dispersion of phage types throughout the host phylogeny, in all but one case, each *E. coli* host harbors only a single prophage. Although phage attachment and infection sometimes depend on O-antigenicity, there is no obvious association between the presence of gokushovirus prophages and particular *E. coli* O-serotypes (**Table S1**). Most gokushovirus prophages were detected in diverse *E. coli* strains isolated from various animals, mainly cattle and marmots, but five prophages were detected in isolates from humans, including one from a urinary tract infection (**Table S1**).

### Reconstituting viable phage from integrated prophages

The integrity of prophage structure and the lack of premature stop codons suggested that they represent intact, functional insertions into bacterial hosts. To confirm this functionality of *Escherichia* gokushovirus prophages, characterize their biology and provide a type strain, we undertook to revive phages from genomic DNA of *Escherichia* strains MOD1-EC2703, MOD1-EC5150, MOD1-EC6098 and MOD1-EC6163, selected to represent the diversity of gokushovirus prophages and hosts (**Figures 1B, C**).

Sequences corresponding prophages from these four *Escherichia* strains were amplified, circularized and transformed into *E. coli* DH5α (**Figure 2A**). Supernatants from the infected DH5α culture were combined with various *E. coli* K12 hosts, resulting in plaques for one of four reconstructed phages (EC6098, derived from *E. marmotae* strain MOD1-EC6098). Occasionally, single colonies in the center of plaques, a bullseye phenotype indicative of lysogeny, were visible (**Figures 2B, C).**

**Figure 2.**
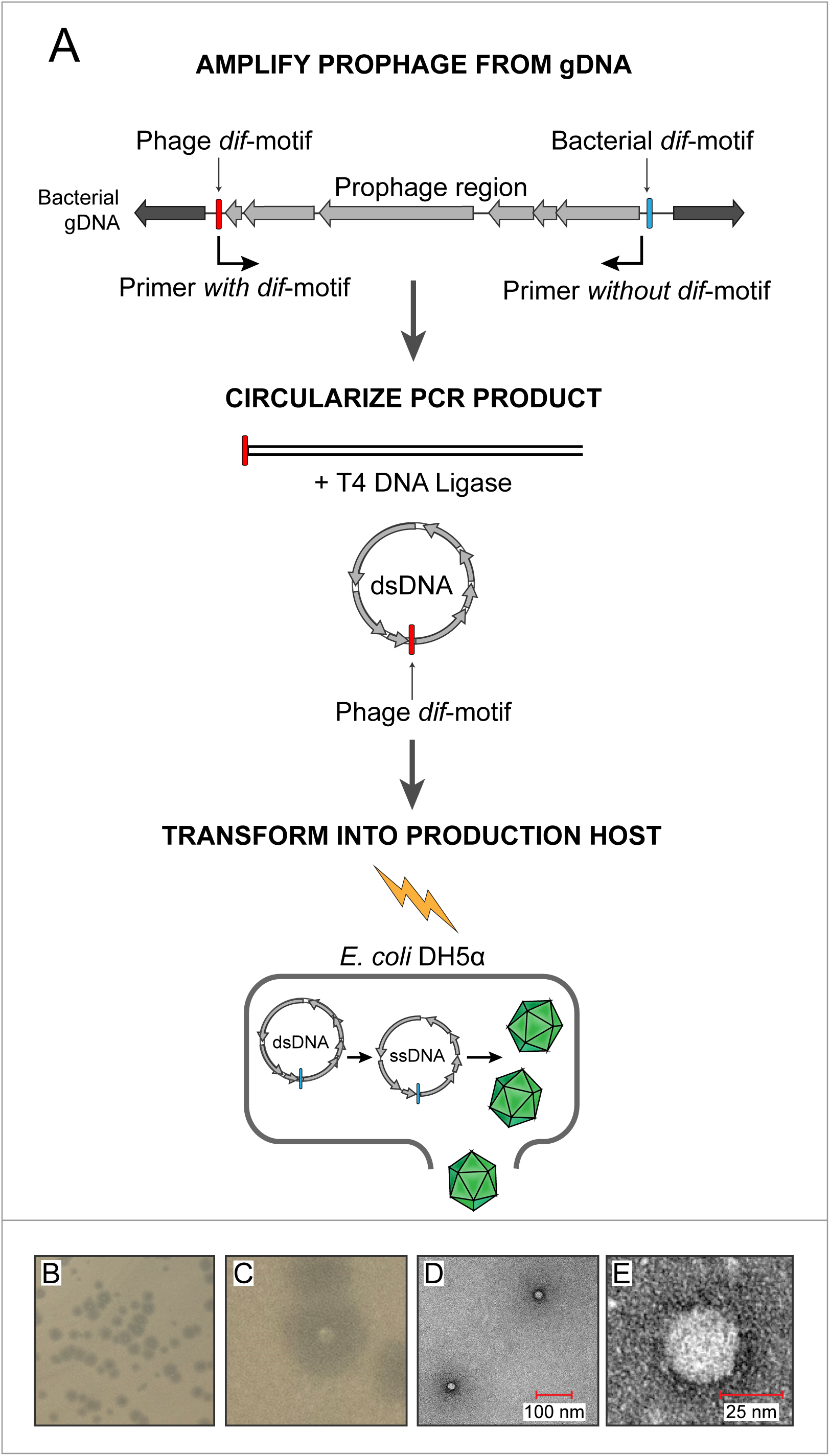
*In vitro* assembly and revival of enterobacterial gokushoviruses. (A) Scheme used to produce viable phage from prophage inserts. The prophage region is amplified using primers (indicated by arrows) that incorporate the phage *dif*-motif but exclude the bacterial *dif*-motif. After circularization of the amplification product, the resulting circular molecule corresponds to the replicative dsDNA form of the phage. Subsequent transformation of the phage into electrocompetent *E. coli* DH5α cells leads to expression of phage proteins, rolling circle replication, and packaging of ssDNA into infective virions. (B) Plaques formed by revived bacteriophage EC6098 after infecting *E. coli* BW25113. (C) Bullseye colony of lysogenized BW25113 situated within a plaque formed by EC6098. (D, E) TEM images of bacteriophage EC6098 viewed at 175,000x and 300,000x magnification.

Electron microscopic observation of pure phage lysates revealed icosahedral virions, 25–30 µm in size and displaying mushroom-like protrusions typical of *Gokushovirinae* (Chipman *et al*., 1998; Diemer and Stedman, 2016) (**Figures 2D, E**). Because *Gokushovirinae* have only been recovered as lytic particles from intracellular bacteria, this represents the first isolation of a gokushovirus able to infect free-living bacteria as well as being able to integrate as a lysogen into bacterial genomes.

### Mechanisms of enterobacterial gokushoviruses integration into host genomes

We next attempted to elucidate the process by which the revived gokushoviruses integrate into the bacterial host chromosome. The presence of circularized phage genomes could readily be detected from bullseye-colonies within plaques, and using primers that flank both sides of the *dif*-motif in host strain BW25113, we recovered products that were enlarged by the length of the phage relative to colonies lacking the prophage (**Figure 3A, B**). Sequencing confirmed that phages integrate downstream of the bacterial *dif-motif*, which remains unchanged.

**Figure 3.**
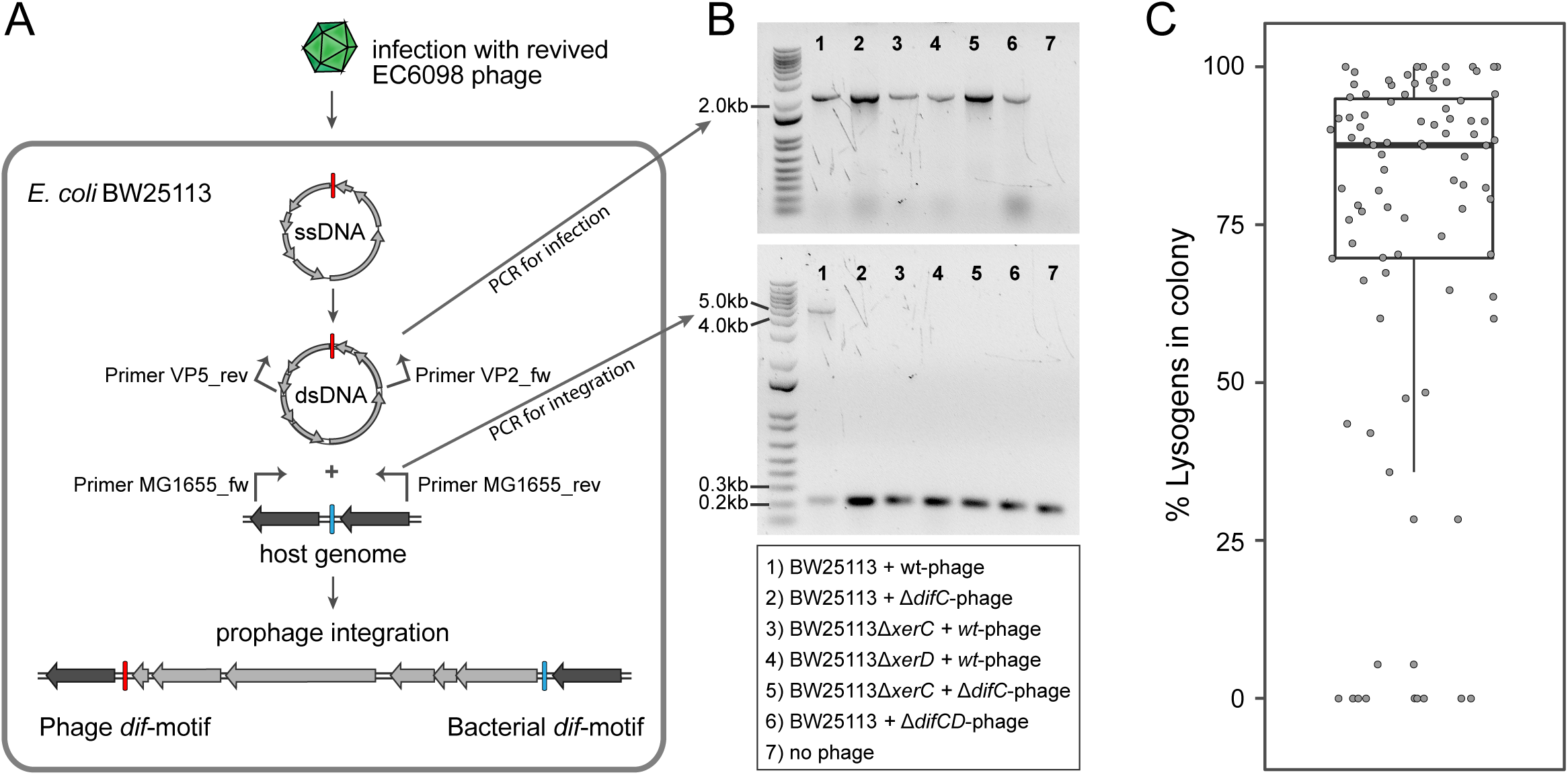
Integration of gokushoviruses into *E. coli* host. (A) Schematic representation of phage integration process as well as detection of circularized EC6098 phage genome and integrated prophages in *E. coli* BW25113. Revived phage EC6098 releases infective ssDNA into host, leading to formation of dsDNA replicative genomes and integration downstream of the bacterial *dif*-motif. Primers VP2_fw and VP5_rev (indicated by arrows on EC6098 genome) anneal to genes flanking the phage *dif*-motif and are used to amplify a ∼2.1-kb product indicative of closed circular phage genomes. Primers MG1655dif_fw and MG1655dif_rev (indicated by arrows on host genome) anneal to sites flanking the bacterial *dif*-motif and amplify either a 210-bp region of bacterial DNA alone or a larger region of integrated prophage flanked by bacterial DNA (∼5 kb). (B) Detection of fragments indicating the presence of circularized (upper gel) and integrated (lower gel) phage from bullseye colonies after infection of BW25113 strains with wild type or mutant phage. (C) Proportion of prophage-carrying cells in clonal lysogenic colonies. Box-and-whiskers plot shows median, 25^th^ and 75^th^ percentiles, and 1.5 inter-quartile range as well as individual datapoints for 87 independently sampled clonal colonies.

Because none of the gokushovirus prophages encodes an integrase, we predicted that host factors XerC and XerD might be responsible for prophage integration, similar to what has been hypothesized for microvirus-prophages in *Bacteroidetes* (Krupovic and Forterre, 2011). Neither Δ*xerC* nor Δ*xerD* mutants of *E. coli* host strain BW25113 resulted in integration, and the Δ*xerC* mutant was restored by complementation with a plasmid expressing the intact version of *xerC*. Similarly, phages with incomplete (*i.e*., lacking either their XerC or XerD binding site) or no *dif-motif*s failed to integrate into host genomes, demonstrating the need for cooperative XerC/XerD binding for successful lysogeny (**Table 1**, **Figure 3B**). However, the retention of *dif-motif*s after integration indicates that this process is reversible: As evident in **Figure 3B**, there is a smaller fragment, in addition to that indicating prophage insertion, that corresponds to those cells in the same colony that do not harbor the integrated prophage. These cells persisted even after multiple rounds of re-streaking, and the median ratio of lysogens to non-lysogenic cells derived from clonal colonies approaches 4:1 (**Figure 3C**). Even in pure cultures, phages are continuously being excised and reintegrated, presumably as a result of XerC/XerD activity. Furthermore, lysogenic strains were immune to infection by EC6098 and, in liquid culture, produced >10^7^ pfu/ml, indicating that integrated prophages result in the production of functional infectious phage particles (**Figure S1**).

**Table 1.** Percentage of lysogenic colonies after phage infection^a^.

| Strain | Phage | Plasmid | % Lysogens |
| --- | --- | --- | --- |
| BW25113 | EC6098 | - | 17.70833333 |
| BW25113Δ*xerC* | EC6098 | - | 0 |
| BW25113Δ*xerC* | EC6098 | pJN105::*xerC* (induced) | 14.58333333 |
| BW25113Δ*xerC* | EC6098 | pJN105::*xerC* (uninduced) | 0 |
| BW25113Δ*xerD* | EC6098 | - | 0 |
| BW25113Δ*xerD* | EC6098 | pJN105::*xerD* (induced) | 0 |
| BW25113Δ*xerD* | EC6098 | pJN105::*xerD* (uninduced) | 0 |
| BW25113 | EC6098Δ*xerC* | - | 0 |
| BW25113 | EC6098Δ*xerD* | - | 0 |
| BW25113 | EC6098Δ*dif* | - | 0 |

<sup>a</sup>Assessed from screening 96 colonies for each strain
<sup>b</sup>Expression induced by addition of 0.1% arabinose

**Figure S1.**
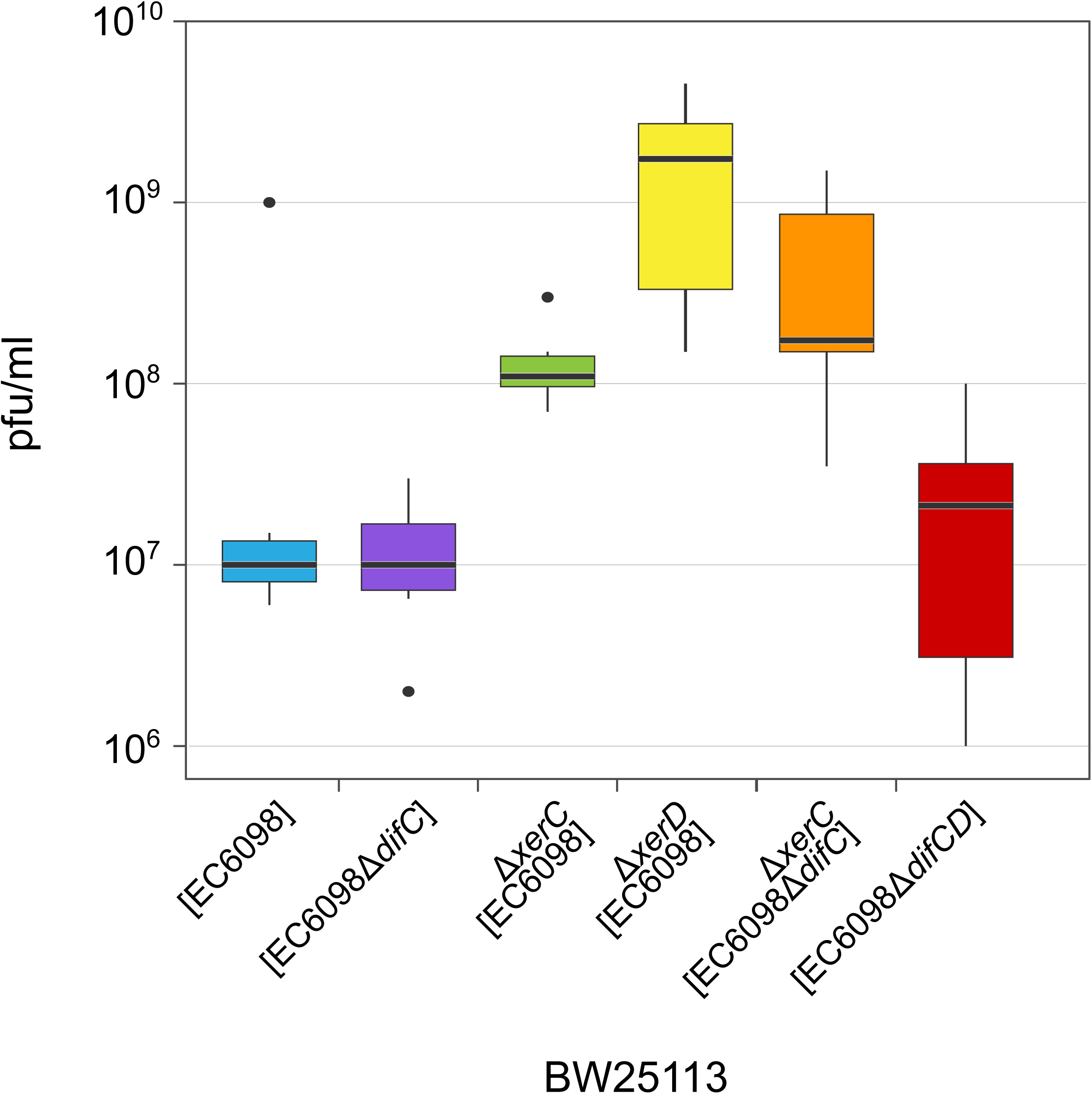
Phage production in the presence and absence of integrated prophages. Plaque-forming units were determined from six (or eight in the case of EC6098Δ*difCD*) independent cultures of BW25113 or BW25113Δ*xerC*, each grown overnight at 37°C from an initial concentration of OD_600_ = 0.7. Box-and-whiskers plots show median, 25^th^ and 75^th^ percentiles (upper and lower hinges) 1.5 inter-quartile range (whiskers). Outliers are shown as individual dots.

### Prophages of enterobacteria form a distinct gokushovirus clade

We initially determined the prevalence of enterobacterial gokushoviruses by interrogating 1699 samples from seven metagenomic studies of human and cattle gut microbiomes for the presence of closely related prophages. In these metagenomes, there were only two prophages corresponding to *E. coli* gokushoviruses, one from the fecal metagenome of an Austrian adult and the other from a Danish infant (**Table S2**).

Phylogenetic analysis of available gokushovirus genomes (*n* = 855; including the enterobacterial prophages discovered in this study, the previously sequenced lytic gokushoviruses from *Chlamydia*, *Spiroplasma* and *Bdellovibrio,* and the metagenomically assembled gokushovirus genomes available from NCBI) returned enterobacterial prophages as a well-supported, monophyletic clade within the *Gokushovirinae* (**Figure 4**). The distinctiveness of this clade, whose members share a conserved gene order and display an average nucleotide identity of >50%, advocates the formation of a new genus within the family *Microviridae* (subfamily *Gokushovirinae*), for which we suggest the name *Enterogokushovirus* on account of a distribution limited to members of the *Enterobactericeae*.

**Figure 4.**
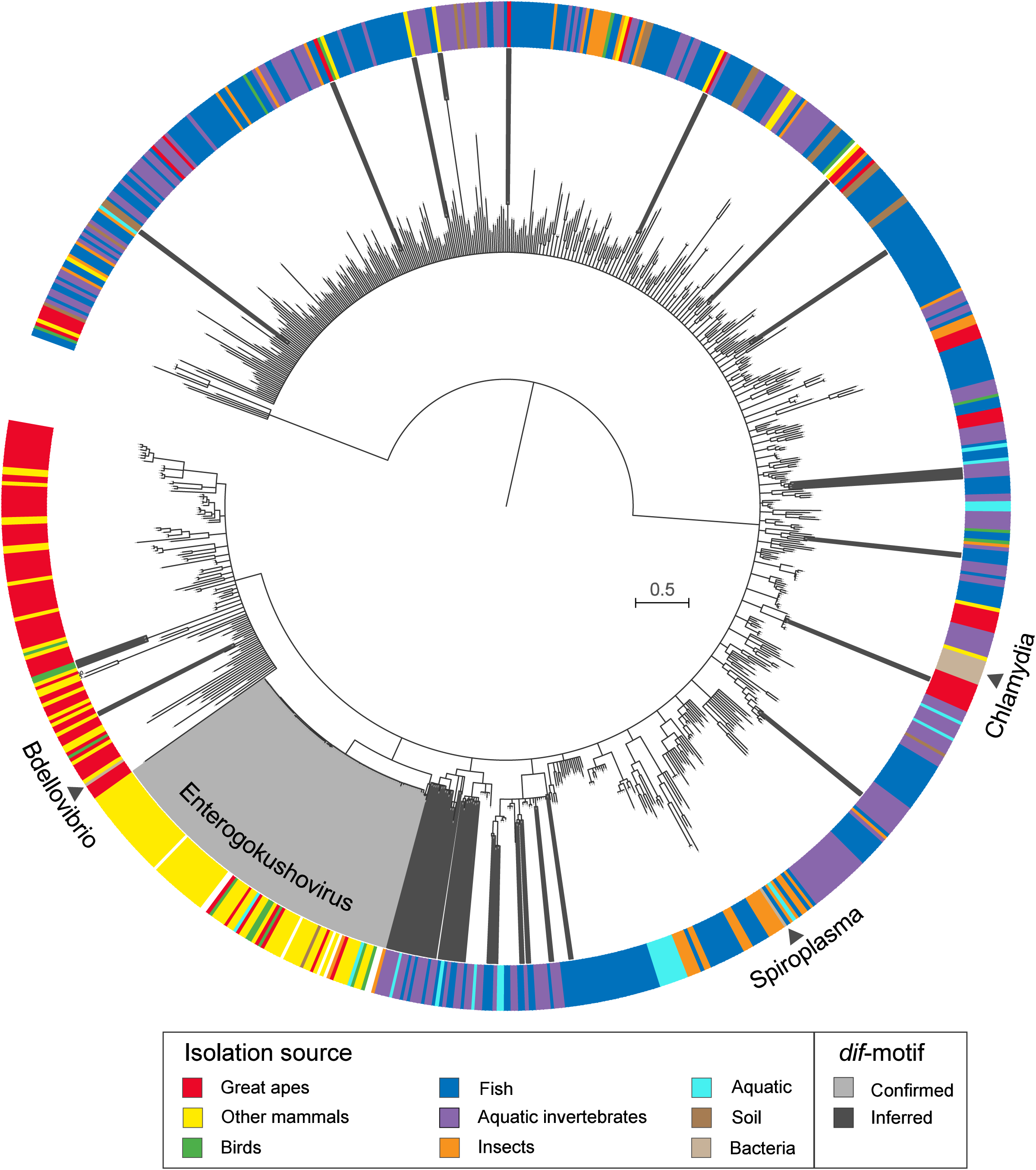
Phylogeny and sources of *Gokushovirinae*. ML tree built from concatenated alignment of VP1 and VP4 protein sequences of 855 gokushovirus genomes. Tree is midpoint rooted, and branch support estimated with 100 bootstrap replicates. Branches with <70% bootstrap support were collapsed. Clades highlighted in light grey indicate *Enterogokushovirus* prophages with recognizable *dif*-motifs; those highlighted in dark grey possess *dif*-motifs identified through an iterative HMM search. Outer ring indicates source of isolation, with black triangles denoting the phylogenetic positions and sources of officially described gokushoviruses. Tree legend corresponds to substitutions per site. Accession numbers for all samples are included in Table S3.

Since a presumably unique feature of enterogokushoviruses among the *Gokushovirinae* is their ability to exist as lysogens, we searched all other gokushoviruses present in metagenomic samples for *dif*-like sequence motifs that might be indicative of lysogenic ability. We detected similar motifs in 48 genomes that were distributed sporadically throughout the gokushovirus phylogeny (**Figure 4**; **Table S3**). The majority of these putative *dif-motif*s occurs in non-coding regions, and those detected within coding regions were typically at the 5’- or 3’-end of a predicted gene, with a nearby alternative start or stop codon that could preserve the genetic integrity of the phage through integration and excision. Aside from the enterogokushoviruses, there are only two other clades in which multiple genomes contain a *dif-*motif. The overall dearth of *dif*-like sequences in gokushoviruses sampled from diverse geographic and ecological settings highlights the distinctiveness of *Enterogokushovirus* genomes and lifestyle.

### Integration into host genomes is not necessary for long-term persistence

The removal of host factors *xerC* or *xerD*, the presence an incomplete phage *dif-motif*, or the lack of a *dif-motif* (as observed in the majority of *Gokushovirinae*) all prevent integration of phages into host genomes (**Table 1**, **Figure 3B**). However, almost all bacterial colonies that survived infections, regardless of whether they were lysogenized or not, were found to contain circular phage DNA (**Figure 3B**). Even cells that lacked any of the factors required for phage integration were both immune to further phage infection and capable of producing infectious particles at levels comparable to those of cultures containing integrated prophages (**Figure S1**).

The absence of alternative integration sites, as verified by inverse PCR, suggested that Gokushoviruses might be able to persist in the host cytoplasm without integration into the host genome. To determine whether this persistence is a transient phenomenon that eventually leads to either the loss of phages or the collapse of bacterial cultures, we performed a month-long serial transfer experiment using a variety of host-phage combinations. With the exception of cultures carrying EC6098Δ*difCD* (*i.e*., those lacking the entire 28-bp *dif-*motif), all bacteria-phage combinations consistently produced phages over the course of this experiment, irrespective of whether they actually contained integrated phages or not (**Figure 5A**). In contrast, 7 of 8 replicate cultures initially producing EC6098Δ*difCD* lost the ability to do so within 3–13 days. The remaining EC6098Δ*difCD* culture produced phage throughout the course of the experiment, albeit at a considerably lower level than all other cultures (**Figure 5B**). Additionally, both lysogenic and non-lysogenic cultures, with the exception of the one producing EC6098Δ*difCD* phages, were immune to infection by further phages (**Figure 5C, D**). Cumulatively, these results indicate that members of the *Gokushovirinae* can survive in lysogenic and lytic states (as exemplified by EC6098 carrying either a complete *dif-*motif or no *dif-*motif at all) or in a pseudolysogenic carrier state (with just one half of an integration site).

**Figure 5.**
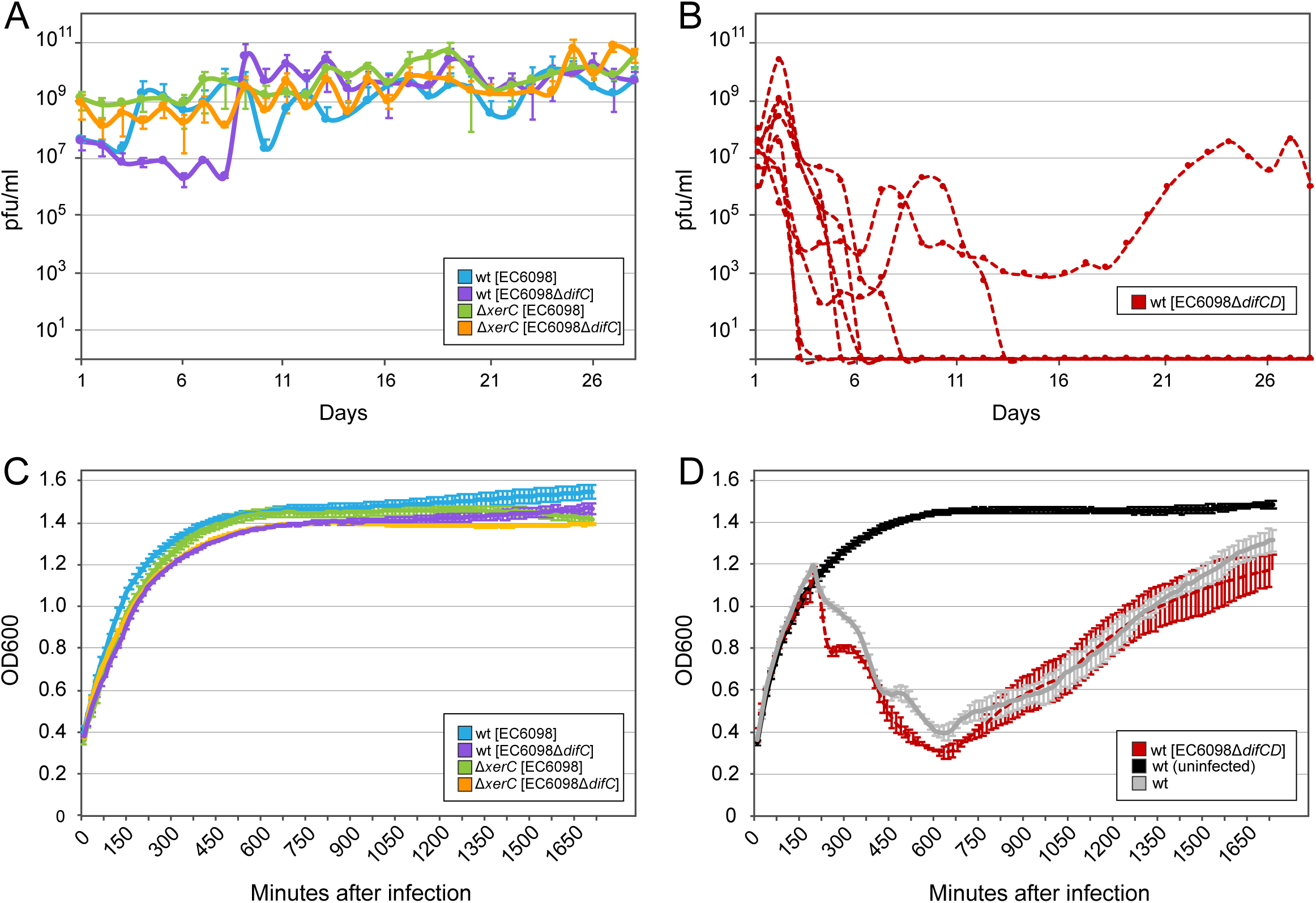
Phage production and susceptibility. (A) Plaque-forming units produced by lysogenic cultures of BW25113 and BW25113Δ*xerC* over the course of 28 daily transfers. (B) Plaque-forming units produced by eight cultures of BW25113 infected with EC6098Δ*difCD* over the course of 28 daily transfers. (C, D) Growth of lysogenic cultures infected with EC6098 after 28 daily transfers. Averages and standard deviations are based on three replicates.

## Discussion

Through the analysis of whole genome sequences and metagenomic databases, we defined a unique genus of *Gokushovirinae* prophages and subsequently synthesized a viable gokushovirus capable of infecting and integrating into the genomes of enteric bacteria. Since *Gokushovirinae* were previously known as exclusively lytic predators of a few intracellular bacteria (King *et al*., 2011), this new genus, *Enterogokushovirus*, offers several new insights into our understanding of this ecologically widespread group of phages. Firstly, by confirming the existence of *Gokushovirinae* in free-living bacteria, we have resolved a seeming paradox in which the most widespread and diverse family of microviruses appeared to be confined to bacterial hosts that are rare pathogens and parasites of eukaryotic cells, such as *Chlamydia* (Wang *et al*. 2019). Moreover, that enteric bacteria regularly serve as hosts for gokushoviruses accounts for their abundance in the healthy human gut microbiome (Manrique and Young, 2017). Secondly, by demonstrating the integration of a revived gokushovirus into the *E. coli* genome, we show that these phages employ survival strategies beyond the lytic infection of hosts.

Experimental characterization of the *Enterogokushovirus* isolated from an environmental strain of *E. marmotae* showed that these phages possess a *dif*-like recognition motif that interacts with the host-encoded recombinases XerC/XerD. In this manner, enterogokushoviruses appropriate a highly conserved bacterial chromosomal concatemer resolution system that enables their integration into host genomes via homologous recombination (Castillo *et al*., 2017). However, phages with only partial DNA-binding motifs exist in a pseudolysogenic state, a condition intermediate between lysogenic and lytic cycles in which circularized phage genomes are present in the cytoplasm and virions are continuously released from the host. This strategy is similar to that observed in CRASSphage (Shkoporov *et al.,* 2018) and other bacteriophages (Siringan *et al*., 2014) and might help explain the stable persistence of microviruses in the human gut (Minot *et al*., 2013).

Despite an exhaustive sampling of gokushoviral diversity from metagenomic datasets, the occurrence of *dif*-positive gokushoviruses is rare outside of the enterogokushoviruses. However, the sporadic distribution of *dif*-motifs among other gokushoviruses indicates that the ability to lysogenize bacterial hosts has been independently gained and lost multiple times during the evolution of this taxon. For example, the exclusively lytic gokushoviruses of *Chlamydia* (Śliva-Dominiak *et al*., 2013) might once have been able to integrate into their hosts’ genomes since extant *Chlamydia* genomes contain coding sequences similar to the gokushoviral replication initiation and minor capsid proteins (Read *et al*., 2000; Rosenwald *et al*., 2014). Perhaps due to the high mutation rate of ssDNA phages, which are typically two orders of magnitude higher than dsDNA phages and estimated to be 10^−5^/nucleotide/day in the human gut (Sanjuan *et al*. 2010, Minot *et al*., 2013), the *de novo* evolution of the XerCD-binding motifs appears to be common, with similar systems existing in other families of phage and mobile-elements (Das *et al*. 2013).

Given the current depths of microbiomes sampling and sequencing, it is surprising that enterogokushoviruses have not previously been identified in human metagenomes. However, their hosts generally do not reach very high abundances in the human gut microbiome, usually constituting less than 0.1% of the total bacterial population (Human Microbiome Project Consortium, 2012). But even among the more than 100,000 sequenced enterobacterial genomes that are currently available, we detected fewer than 100 gokushovirus prophages, and no gokushovirus prophages were detected in the sequenced genomes of other bacteria. This rarity of gokushoviruses existing as prophages might result from the process by which they integrate into genomes: although possession of a *dif*-motif provides a simple, passive way for phages to integrate into host genomes, they also facilitate prophage excision. In fact, the XerCD-mediated removal of genetic elements has attained application in molecular biology (Bloor and Cranenburgh, 2006). For example, a *Vibrio cholerae* phage, CTX secures itself as a prophage by destroying the *dif* site upon insertion, (Val *et al*., 2005), whereas *Vibrio* prophages that retain their *dif-motif*s are only rarely detected in sequenced genomes (Das *et al*., 2011). The abundance of gokushoviruses in the environment, as apparent from their regularity in metagenomic datasets, suggests that they are only transient residents of bacterial chromosomes and usually occur in pseudolysogenic or lytic states in the wild.

The presence of numerous and diverse gokushovirus prophages in a wide variety of *E. coli* strains now makes it possible to elucidate aspects of gokushovirus biology in a comparative evolutionary framework. For example, the role of the ∼300-bp hypervariable region in the major capsid protein, which varies considerably among enterogokushoviruses and among *Gokushovirinae* in general (Roux *et al*. 2012, Hopkins *et al.,* 2014), has been the subject of some speculation. This region forms the characteristic protrusions at the three-fold axis of symmetry in *Spiroplasma*-microvirus virions and has long been hypothesized to be involved in the host range determination (Chipman *et al*., 1998). Of the four synthesized genomes employed in this study, we observed that only one (EC6098) produced viroids capable of infecting *E. coli* K12. That each genome displayed a unique hypervariable region in its VP1 gene suggests that host range determinants are linked to these different capsid regions.

The discovery of this group of phages offers several new directions for the study of the *Microviridae*. Demonstrating that the host ranges of the *Gokushovirinae* extends beyond intracellular bacteria increases the likelihood that additional members of this prevalent group of phages will be isolated from appropriate hosts. Furthermore, the amplification of entire ssDNA phage genomes and subsequent transformation into appropriate hosts, as demonstrated in this study and prior with *de novo* synthesized *phiX* (Smith *et al*., 2003) can aid in the description of other *Microviridae* subfamilies. A promising target for this would be the *Alpavirinae*, which have been detected as prophages of *Bacteroidetes* but so far remained without isolates, and thus been denied official recognition (Roux *et al.,* 2012). In conclusion, the value of isolating enterogokushoviruses goes beyond their abundance in nature and provides a genetically manipulable model system that will further the understanding of this group as a whole (Shkoporov and Hill, 2019).

## Acknowledgements

The authors thank the Federal Department of Agriculture for providing *Escherichia spp.* strains and DNA, Steven Kyle for assistance in experiments, Dwight Romanovicz for assistance with electron microscopy, and Kim Hammond for figure design. This study was funded by NIH award R35GM118038 to H.O.

## Author contributions

PCK planned and executed the study, and performed all experiments and analyses. PCK and HO wrote and edited the manuscript.

## Competing Interests

The authors declare no competing interests.

## Data availability statement

The authors confirm that the data used in this publication (accession numbers, sequences and alignments) are available in the paper, its supplementary files and/or from the authors upon request.

## Material & Methods

### Detection and phylogeny of Microvirus prophages and their hosts

Using blastp, we queried the NCBI nr database (April 2019) with the *Chlamydia* Phage 4 major capsid protein VP1 (NCBI Gene ID 3703676) and discovered several sequences with E-values less than 10E-4 within the *Enterobacteriacae*. We used these hits to query the genomes of all *Enterobactericaea* in the NCBI microbial draft genome database (April 2019) using blastn and downloaded the complete genome sequences of strains returning hits with e-values less than 10E-4 **(Table S1).** Bacterial chromosome contigs containing the VP1 gene were visually inspected in Geneious R9 (www.geneious.com) for the presence of prophage insertion boundaries by searching for identical 17-bp sequences within the 5-kb regions both up- and downstream of the VP1 gene. Prophage genes were annotated with GLIMMER3 (Delcher *et al*., 2007) using default settings, specifying a minimum gene length of 110 bp and a maximum overlap of 50 bp. Initial alignments of prophage regions were made with ClustalO 1.2.4 (Sievers *et al*., 2011), which were refined manually to accommodate hypervariable regions and the phage insertion sites at the 3’ and 5’ ends of the alignment. We then built a maximum likelihood phylogenetic tree of enterobacterial prophages with RAxML 8.0.26 (Stamatakis, 2014) using the GTR+GAMMA substitution model and 100 fast-bootstrap replicates, and visualized the resulting tree in FigTree 1.4.3 (http://tree.bio.ed.ac.uk/software/figtree/).

To evaluate the distribution of prophage hosts within the broad diversity of *E. coli* at large, we produced core genome alignments of prophage hosts and representative genomes from the ECOR collection (**Tables S1** and **S2**) based on protein families satisfying a 30% amino-acid identity cutoff (USEARCH 11, Edgar, 2010), which were aligned with MUSCLE 3.8.31 (Edgar, 2004), as implemented in the BPGA 1.3 pipeline (Chaudhari, 2016). The maximum likelihood phylogenetic tree of core genome alignments was built with IQTree 1.6.2 (Nguyen, 2015), using the JTT substitution model and 100 bootstrap replicates.

### Recovering prophages from metagenomes

To assemble prophages from metagenomic datasets, we downloaded SRA files from seven BioProjects (PRJEB29491, PRJNA362629, PRJNA290380, PRJNA352475, PRJEB6456, PRJNA385126 and PRJEB7774) and performed initial trimming and quality filtering with BBDuk (Bushnell, 2014a) with options ktrim=r k=23 mink=11 hdist=1 tbe tbo. Reads having a minimum nucleotide sequence identity of 50% to sequences of enterobacterial prophages (above) as determined by BBMap (Bushnell, 2014b) were assembled into contigs using MEGAHIT 1.1.3 (Li *et al.,* 2015) implemented with default settings, and contigs >1000 bp were retained.

### Phylogenetic analysis of Gokushovirinae

We downloaded a total of 1284 metagenome-assembled genomes (MAGs) of microviruses (**Table S3**), which were then reannotated in GLIMMER3 (Delcher *et al*., 2007) using default settings, with a minimum gene length of 110 bp and a maximum overlap of 50 bp. We recovered homologues to the conserved major capsid protein VP1 and replication initiation protein VP4 in the set of metagenome-assembled microviruses using PSI-BLAST searches and querying with VP1 and VP4 proteins from detected enterobacterial gokushoviruses, gokushovirus genomes of *Chlamydia*, *Spiroplasma* and *Bdellvibrio*, and *Bullavirinae* phage *phiX*174. After individual protein alignments using Clustal Omega 1.2.4, we concatenated the VP1 and VP4 alignments, and removed all sites with >10% gaps to decrease the amount of spuriously aligned sites using Geneious R9. The initial phylogenetic tree of all microviruses was built with IQTree 1.6.2 using the LG+F+R10 substitution model as determined by ModelFinder (Kalyaanamoorthy *et al*., 2017), and branch support was tested using 1000 ultra-fast bootstrap replicates (Hoang *et al*., 2018) and 1000 SH-aLRT tests. Collapsing all branches with <95% bootstrap support and <80% SH-aLRT support yielded a single, well-supported clade containing all known *Gokushovirinae*, and all subsequent alignments and phylogenetic trees were refined by including only those genomes represented in this clade, with branch support assessed using 100 standard bootstrap replicates.

### Identification of *dif*-motifs in gokushovirus MAGs

To search for *dif*-motifs in enterobacterial prophages, we first performed an alignment of all enterobacterial prophage *dif-motif* sequences in the curated set of bacterial *dif*-motifs from Komo *et al*. (2011) using Clustal Omega 1.2.4. We used the resulting alignment to build a Hidden-Markov-Model using hmmer 3.2.1 (Wheeler *et al.,* 2013) and performed an iterative search for *dif*-like motifs in all gokushovirus-like MAGs. Due to the variation in phage and bacterial *dif*-motifs, the variation in these motifs among bacteria, and the short length of the target sites, only confirmed *dif*-motifs of enterobacterial prophages reached the E-value cutoffs of 10E-4. A large number of hits fell below of this threshold and were treated as potential *dif*-motifs if they possessed at least 15 bp identical to confirmed *dif*-motifs and occurred in the short non-coding regions of MAGs. All hits within coding regions were removed as likely representing false positives, with the exception of those within the N-terminus of VP4 (as occasionally observed in *Escherichia* gokushovirus prophages).

### Resurrection and modification of prophages

DNA fragments representing gokushovirus prophages were amplified from *E. coli and E. marmotae* strains with Phusion polymerase (NEB) from 10 ng of genomic DNA using primer pairs listed in **Table S4** and under the following PCR conditions: 98°C for 3 min; 30 cycles of 98°C for 15 sec, 50°C for 15 sec, 72°C 2:30 min; followed by 72°C 10 min. Amplified fragments of ∼4.5 kb corresponding to gokushovirus prophages were purified from agarose gels using the Monarch DNA Gel Extraction Kit (NEB) and eluted in 20 µl ddH_2_O. Blunt ends of the purified linear fragment were phosphorylated with T4 Polynucleotide Kinase (ThermoFisher) followed by overnight treatment with T4 DNA Ligase (NEB) to form circular genomes. Ligation mixtures were heat-inactivated, desalted, transformed into *E. coli* DH5α and incubated for 1 hr in 1 ml SOC medium at 37°C. After this recovery period, cultures were grown overnight in 5 ml of LB medium at 37°C with mild shaking (200 rpm). Viable bacteriophages were harvested by centrifuging the culture for 5 min at 5000 g to pellet bacterial cells and then by filtering the supernatant through 0.45 µm syringe filters. The presence and identity of phages were confirmed through standard spot assays (see below) and nucleotide sequencing (see **Table S4**).

### Plasmid construction and complementation of knockout mutants

To construct complementation plasmids, we first amplified the *xerC* and *xerD* genes from *Escherichia* coli BW25113 with primers XerC_fw_EcoRI and XerC_rev _SacI or XerD_fw_EcoRI and XerD_rev_SacI (**Table S4**) under the conditions listed above. PCR products were purified using the Monarch DNA Gel Extraction Kit (NEB) and eluted in 20 µl ddH_2_O. PCR products and expression plasmid pJN105 (Newman and Fuqua, 1999) were digested with *Eco*RI and *Sac*I (NEB) for 37°C for 1 hr, followed by heat inactivation for 10 min at 80°C and overnight ligation at a 1:3 vector-to-insert ratio using T4 DNA Ligase (NEB) at 4°C. One microliter of ligation mixtures were transformed into electrocompetent BW25113Δ*xerC or* Δ*xerD* mutants, and transformants were selected for growth on LB agar plates supplemented with 10µg/ml gentamycin.

### Phage and bacterial culture

Environmental *Escherichia* strains MOD1-EC2703, MOD1-EC5150, MOD1-EC6098 and MOD1-EC6163, *Escherichia coli* K12 derivates MG1655, DH5alpha and BW25113, and KEIO collection strains BW25113Δ*xerC* (KEIO Strain JW3784-1) *and* BW25133Δ*xerD* (KEIO Strain JW2862-1) were grown at 37°C in LB liquid media (supplemented with of 50 µg/ml kanamycin for the KEIO knockout strains). Expression of *xerC* or *xerD* genes in BW25113Δ*xerC* and BW25133Δ*xerD* containing pJN::xerC or pJN::xerD was induced by addition of 0.1% arabinose.

To prepare agar-overlays, cells from 100 µl of overnight culture were pelleted, resuspended in PBS, combined with 100 µl of phage and incubated at room temperature for 5 minutes prior to addition 3 ml of 0.6% LB-agarose and plating onto LB agar. To increase the phage concentrations, we harvested phage lysates from plates exhibiting confluent lysis after overnight growth at 37°C. Phage titers were determined by spotting dilutions of lysates onto agar-overlay plates with 100 µl of overnight cultures of host strains and incubating plates overnight at 37°C. Liquid-infection assays were performed in 96-well plates by adding 2 µl of phage lysate (∼10^8^ pfu/ml) to 200 µl of overnight culture diluted with LB to OD_600_ = 0.4 and measuring growth and lysis at 37°C with 200 rpm shaking at 15-min intervals on a Tecan Spark 10M plate reader.

Serial transfers experiments were performed by inoculating lysogenic colonies in LB, diluting overnight cultures to OD_600_ = 0.7, and then transferring 2 µl of the diluted culture into 2 ml of LB medium. After 18–24 hours incubation at 37°C with shaking, 2 µl of culture was transferred to 2 ml of fresh LB. This process was repeated for 28 days, and each day, phages were titered in the manner described above.

### Detection of circularized phages and prophages

The presence of lysogens and circularized phage was determined by PCR assays of liquid overnight cultures from bullseye colonies derived from a single phage infection using primers MG1655_fw and MG1655_rev, which flank the bacterial *dif*-motif, and VP2_fw and VP5_rev, which anneal up- and downstream the phage *dif*-motif in circularized phage genomes (**Table S4**). Ratios of lysogenic to non-lysogenic cells in individual cultures were measured by re-streaking single bullseye colonies three times, selecting and resuspending a single resulting colony in LB, and re-plating it onto LB-agar plates. Colonies were grown in liquid culture overnight and assayed by PCR with primers MG1655_fw and MG1655_rev to detect prophage integration, as described above. PCR products were resolved on 1% agarose gels, and the intensity of the PCR products representing integrated prophage and non-integrated sites was measured with ImageJ 1.52a (http://imagej.nih.gov/ij).

Prophage integration sites were confirmed by inverse PCR (Ochman *et al.,* 1988) as follows: 10 ng of DNA derived from colonies of BW25113 and BW25113Δ*xerC* infected with either EC6098 wild type or EC6098Δ*difC* were cut with restriction enzyme *Hin*dIII (NEB) for 1h at 37°C. Reactions were heat inactivated at 80°C for 10 minutes, and then circularized with T4 DNA Ligase (NEB) overnight at 4°C. Primer VP2_rev, binding the 3’-end of VP2 and facing upstream, and primers Circle_1 and Circle_2, which both bind in conserved regions downstream of VP2 and face downstream, were used to amplify circularized ligation products using Phusion polymerase (NEB) under the following PCR conditions described above. PCR products were resolved on 1% agarose gels, and all detected bands were extracted and sequenced. Phage integration was confirmed when single-read sequences contained both phage and bacterial DNA.

### Electron microscopy

Two milliliiters of high-titer phage lysate resuspended in PBS was layered on top of a CsCl step gradient (2 ml each of p1.6 to p1.2 in PBS) and centrifuged in a Beckman Coulter Optima L-100k Ultracentrifuge at 24,000 rpm for four hours. After centrifugation, fractions were collected in 0.5ml steps by puncturing the centrifuge tube at the bottom. To determine which fractions contained phage, PCR was performed using primers nocode_fw and VP2_rev as described above, using 1 µl of each fraction as template. Fractions containing phage were desalted using an Amicon Ultra-2ml Ultracel-30k filter unit and resuspended in water. For electron microscopy, viral suspensions were pipetted onto carbon-coated grids, negatively stained with 2% uranyl acetate, and imaged with a Tecnai BioTwin TEM operated at 80kV.

## Supplemental Information Titles and Legend

**Table S1. Enterobacteria harboring gokushovirus prophages.**

**Table S2. Reference strains for host-phylogeny.**

**Table S3. Gokushovirus genomes.**

**Table S4. Primers used in this study.**

